# Predicting S. aureus antimicrobial resistance with interpretable genomic space maps

**DOI:** 10.1101/2023.02.24.529878

**Authors:** Karina Pikalyova, Alexey Orlov, Dragos Horvath, Gilles Marcou, Alexandre Varnek

## Abstract

Increasing antimicrobial resistance (AMR) represents a global healthcare threat. Methods for rapid selection of optimal antibiotic treatment are urgently needed to decrease the spread of AMR and associated mortality. The use of machine learning (ML) techniques based on genomic data to predict resistance phenotypes serves as a solution for the acceleration of the clinical response prior to phenotypic testing. Nonetheless, many existing ML methods lack interpretability and do not implicitly incorporate visualization of the sequence space that can be useful for extracting insightful patterns from genomic data. Herein, we present a methodology for AMR prediction and visualization of sequence space based on the non-linear dimensionality reduction method □ generative topographic mapping (GTM). This approach applied to data on AMR of >5000 S. aureus isolates retrieved from the PATRIC database yielded GTM models with reasonable accuracy for all drugs (balanced accuracy values ≥0.75). The GTMs represent data in the form of illustrative 2D maps of the genomic space and allow for antibiotic-wise comparison of resistance phenotypes. In addition to that, the maps were found to be useful for the analysis of genetic determinants responsible for drug resistance based on the data from the PATRIC database. Overall, the GTM-based methodology is a useful tool for the illustrative exploration of the genomic sequence space and modelling AMR and can be used as a tool complementary to the existing ML methods for AMR prediction.

**Availability:** https://doi.org/10.5281/zenodo.7101559

## 1 Introduction

**T**HE discovery of antibiotics marked the revolution in bacterial infection treatment that led to a significant decrease in associated mortality and morbidity. However, the ability of pathogenic bacteria species to adapt and resist the treatment on par with inappropriate practice of antibiotic usage lead to an increase in the number of deaths from bacterial infections. According to recent research, more than 1 million deaths can be attributed to antimicrobial resistance (AMR) in 2019 [1]. Until immediate measures are taken, it is estimated that the death toll from AMR could soar to 10 million lives per year by 2050 [2]. One of the important measures that can be taken to slow down the spread of AMR is the optimization of antimicrobial drugs’ use and stewardship [3].

To establish a diagnosis and prescribe proper treatment for patients with bacterial infection, healthcare specialists usually rely on *in vitro* antimicrobial susceptibility testing (AST) [2]. AST is considered to be a ‘golden standard’ for choosing optimal antibiotics to which the pathogen in question is susceptible [4]. However, AST is a time-consuming and labour-intensive process [5]. Hence, the development of rapid front-line diagnostics tools for better clinical decision-making is urgently needed [2].

Due to increased availability and rapidity, genotypic methods represent a cost-effective alternative to phenotypic AST methods. Genotypic methods consist in the sequencing of the pathogenic organism and subsequent application of rule- or ML-based approaches for resistance predictions. The rule-based approaches predict AMR on the basis of preliminary knowledge of AMR genes or resistance-inducing mutation patterns. Hence, the performance of such approaches is especially good for well-studied organisms [6]. Machine learning (ML) methods, on the other hand, correlate genomic features with phenotype; they are trained on raw genomic data associated with experimentally measured AMR phenotypes. In recent years, numerous approaches for ML-based antibiogram predictions were suggested [7]. Various ML algorithms including AdaBoost [8], Random Forest (RF) [9], [10], Support Vector Machine (SVM) [11], CATBoost [9], Neural Networks (NNs) [10], [12] were applied to predict resistance/susceptibility labels or minimum inhibitory concentration (MIC) values of antimicrobial agents for bacterial isolates. The genomic data used as an input to these algorithms can be represented in numerous ways: as whole-genome sequences, genome contigs, AMR genes, large-scale pangenome data, short sequencing reads, etc. It is usually encoded by the means of alignment-based and alignment-free methods, e.g. sets of single nucleotide polymorphisms (SNPs) and k-mers respectively [13]. For a detailed description of the approaches, the reader is referred to the recent review by [7].

Besides high precision, the interpretability of ML plays an important role in the incorporation and adoption of the algorithm into the clinical diagnostics pipeline. To address the issue of clinical utility, several approaches to interpretable ML methods for AMR prediction were suggested [14], [15]. For example, some of the current interpretability-focused approaches rely on rule-based learning algorithms that are combined with sample compression theory [14] or lasso-penalized logistic regression model [15]. Numerous ML tools were also suggested for genomic data visualization and analytics, namely, dimensionality reduction techniques such as t-distributed stochastic neighbour embedding (t-SNE) [16], generative topographic mapping (GTM) [16], stochastic cluster embedding (SCE) implemented in a Mandrake tool [17], principal components analysis (PCA) [18], [19], uniform manifold approximation and projection (UMAP) [18], topology data analysis methods [20], [21], etc. These methods enable gaining insight on population structure via illustrative exploration of the genomic space.

Ideally, methods should provide both data visualization and performance guarantees. For this purpose, we suggest a methodology for the analysis of bacteria genomic data and predicting AMR with emphasis on the model’s explainability and examination of markers of resistance. The methodology is based on the non-linear dimensionality reduction method – Generative Topographic Mapping (GTM) [22], [23]. Previously, the GTM was shown to be effective for the analysis of large-scale chemical [24] and biological data [16], [25], [26]. Recently, GTM was found to be particularly effective in predicting HIV drug resistance, visualization of sequence space of HIV proteins and analysis of the HIV isolates’ mutation patterns [25].

Herein, the GTM was applied for genomic space visualization and AMR predictions for *Staphylococcus aureus* (*S. aureus*), one of the six most pathogenic bacteria species ESKAPE (*Enterococcus faecium, Staphylococcus aureus, Klebsiella pneumoniae, Acinetobacter baumannii, Pseudomonas aeruginosa* and *Enterobacter* spp.) recognized by WHO. The uniqueness of the herein presented GTM-based methodology lies in its simultaneous ability of visualization of sequence space and prediction of antibiotic resistance given data on AMR profiles and genetic determinants. Albeit the performance of the GTM was lower as compared to other state-of-the-art ML methods, the GTM-based models appeared to be useful for predictions (BA values • 0.75 for all antibiotics) and illustrative analysis of *S.aureus* genomic space in the context of the resistance.

## 2 Methods

### 2.1 Data acquisition and pre-processing

Pathosystems Resource Integration Center (PATRIC) database [27] was used to retrieve bacterial genomes of *S. aureus* with available AMR profiles, e.g. labelled as resistant or susceptible according to experimental minimal inhibitory concentration (MIC) breakpoints. A contigs-based representation thereof was downloaded from the PATRIC database. The detailed pre-processing protocol is provided in Supplementary Information Text S1. The final number of pre-processed *S. aureus* isolates was 5773. The overall number of antibiotics after pre-processing was 11. The antibiotics against *S. aureus* and the corresponding number of resistant and susceptible data points extracted from the PATRIC database after decreasing the majority class size, i.e. by randomly removing data points of the majority class until reaching the class ratio of 1:1 are reported in Table S1.

### 2.2 Descriptors

Two commonly used approaches for genomic information representation are single nucleotide polymorphism (SNPs) matrices and k-mer profiling. The former approach, capturing alterations of base pairs in a DNA among collections of aligned sequences, is multiple sequence alignment (MSA)-dependent, is computationally costly and can be error-prone due to lateral gene transfer events frequently encountered in bacteria [14]. The latter approach, being alignment-free, describes each genome by the subsequences of nucleotides of a defined length (k-mers) that are present within a given DNA sequence. In this way, sequences can be described by counting the number of times each k-mer occurs in a sequence or k-mer presence/absence profile within the sequence, thus avoiding computationally expensive MSA [14]. Furthermore, quivalent k-mers can be combined into “representative k-mers” denoted as unitigs. In this work, alignment-free approaches were used to represent bacterial genomes, namely k-mers’ counts and unitigs were computed. The procedure used for the generation of unitigs is explained in Supplementary Information Text S2 and Fig. S3.

### 2.3 Generation of k-mers

Given the lengths of bacterial genomes, the number of generated k-mers can soar up to tens of millions, depending on k-mer length. Such datasets with an extremely small number of samples relative to the number of features, commonly denoted as ‘fat data’, are reputedly difficult ML challenges [14]. Therefore, an additional preprocessing procedure was applied to reduce the number of features. Namely, PCA and kernel PCA (kPCA) were used for feature space reduction. In more detail, k-mers (10-, 15-, and 30-mers) were generated using two programs: dsk for 10-mers [28] and Kmer-db for 10-, 15-, and 30-mers [29]. Herein, Kmer-db was used to calculate k-mer counts and estimate distances between sequences using different metrics (i.e. Jaccard, cosine, max, min, ani, Mash). The Kmer-db approach was applied for the generation of kernel-PCA pre-processed descriptors, whereas for PCA pre-processed descriptors the dsk tool was used since it enabled direct extraction of k-mers’ counts in the text format. More details on libraries used for PCA and kPCA, as well as a list of all kernels tried herein are given in Supplementary Information Text S3.

### 2.4 Generative Topographic Mapping

Generative Topographic Mapping (GTM) first proposed by Bishop [22] is a non-linear dimensionality reduction technique. The GTM principle consists in the insertion of a flexible hypersurface denoted « manifold » into the high-dimensional descriptor space to attain the maximum coverage of densely populated zones of the sequence space. Once the optimal manifold geometry allowing to encompass as many items from the frame set (a set of representative genomes taken for the study) as possible is found, the points can be projected onto it. The manifold is defined by a set of Gaussian Radial Basis Functions (RBFs). The RBF is used as the centre of a gaussian distribution that serves to estimate the likelihood of a sequence to be projected onto the manifold, and its responsibility (probability of a sequence to be assigned to a given node). Hence, through responsibilities, each sequence in the multi-dimensional space is associated with nodes of the manifold [24]. The geometric centre of the responsibilities of a given sequence defines its position on the manifold – the latent space. The major GTM advantage over other non-linear dimensionality reduction methods such as Kohonen Self-Organizing Maps (SOM) [30] is its « fuzzy logics ». An item’s projection is a probability (responsibility) vector. Namely, the sequence is not located in a particular node, but associated to several nodes, with varying “responsibilities” that sum up to 1.0. Such projection of items allows for the creation of GTM landscapes locally coloured by mean values of properties (density or drug resistance). Multiple landscapes can be accommodated on one manifold, i.e. one sole manifold can be coloured by resistance values to different drugs, hence generating distinct resistance landscapes for each of the antibiotics on a same map. GTM possesses four hyperparameters: map resolution, number of hidden Radial Basis Functions (RBF), regularization coefficient and width of an RBF that can be optimized all along with descriptor sets. The optimization of the former and latter is usually done by the means of Genetic Algorithm (GA). More details on the GTM algorithm are given in Supplementary Information Text S4.

There are several reasons for using GTM to visualize bacterial population structure and attempt antibiotic resistance predictions. Topological methods are believed to be especially useful for bacterial genomic data analysis and visualization. Events commonly occurring in bacteria such as recombination and horizontal gene transfer complicate the sequence analysis by tree-based algorithms such as phylogenetic trees [21]. The advantage of the GTM, as opposed to other non-linear dimensionality reduction methods such as t-SNE [31] relies in its reproducibility and ability to update the model with new data, without retraining of the manifold [23]. In addition to that, its fuzzy-logical nature allows for capturing subtle differences between sequences by modulating sequence node responsibilities, as opposed to self-organizing maps (SOM), where subtle variations of an item would likely not alter its unique node of residence [24], [30].

### 2.5 Genetic algorithm

GA is a stochastic evolutionary algorithm allowing one to choose the best set of GTM hyperparameters and descriptors [32]. Herein, the GA was used for Pareto-front driven optimization of BA values for a set of drug-specific models. The quality of the GTM (i.e. predictive model) with the given hyperparameters and descriptors is estimated by its predictive propensity. Herein, the generated landscapes were evaluated by the averaged balanced accuracy (BA) (average proportion of sequences predicted correctly for each class: susceptible and resistant) for each drug in a 5-fold cross-validation procedure repeated 5 times.

### 2.6 Assessment of prediction accuracy of models

To assess the predictive accuracy of the models, the fivefold cross-validated averaged balanced accuracy was estimated upon 5 repeats:

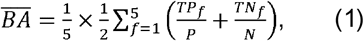

*TPf* - the number of truly resistant sequences predicted as resistant in the fold f, *TNf* - the number of truly susceptible sequences predicted as susceptible in the fold f, *N*-the total number of sequences actually be-longing to the susceptible class, *P-* the total number of sequences actually belonging to the resistant class.

In more detail, herein, the drug-specific resistance landscapes were created using one common manifold. Each of the landscapes represents a classical single-task model aimed to predict the property (resistance) of genomes. Since each of the drug-specific landscapes possesses the same manifold hyperparameters, the quality of the landscape-based models relies on the inherent characteristics of the common manifold they are built on [33]. The maps were considered as optimal if the set of GTM hyperparameters along with descriptors’ sets allowed for consensus maximization of BA values of drug-specific landscape-based models.

In summary, the hyperparameters for manifold describing *S.aureus* sequence space were selected during cross-validation with respect to the mean predictive performance (BA values) of *S.aureus* isolates drug resistance toward nine antibiotics. Two more drugs (oxacillin and sulfamethoxazole/trimethoprim combination antibiotic (SMZ-TMP)) were chosen to validate the ability of the manifold to host predictive resistance landscapes for drugs not used during map fitting process and ensure that the model’s performance is preserved.

### 2.7 Comparison to other machine learning methods

GTM-driven antibiotic resistance predictive ability was compared with other state-of-the-art ML algorithms such as RF [34], SVM [35] and Gradient boosting (GB) [36] implemented in the scikit-learn library (v. 1.1.1) [37]. The hyperparameters tuned during optimization by grid search are reported in Supplementary Information Text S5. Evaluation of the model performance was made using 5-fold cross-validation repeated 5 times (Equation 1).

## 3 Results and Discussion

### 3.1 Benchmarking of GTM and state-of-the-art ML methods

The benchmarking of the GTM-based multi-task models (models optimized simultaneously for 9 antibiotics) with state-of-the-art ML methods such as RF, GB, and SVM in single-task mode display higher performance of latter methods in comparison with GTM upon 5-fold crossvalidated balanced accuracy evaluations repeated 5 times (Fig. 1, Fig. S10). The accuracies of the herein presented ML models and existing AdaBoost classifiers [8] are similar in the case of methicillin resistance predictions for *S. aureus* isolates.

**Fig. 1.**
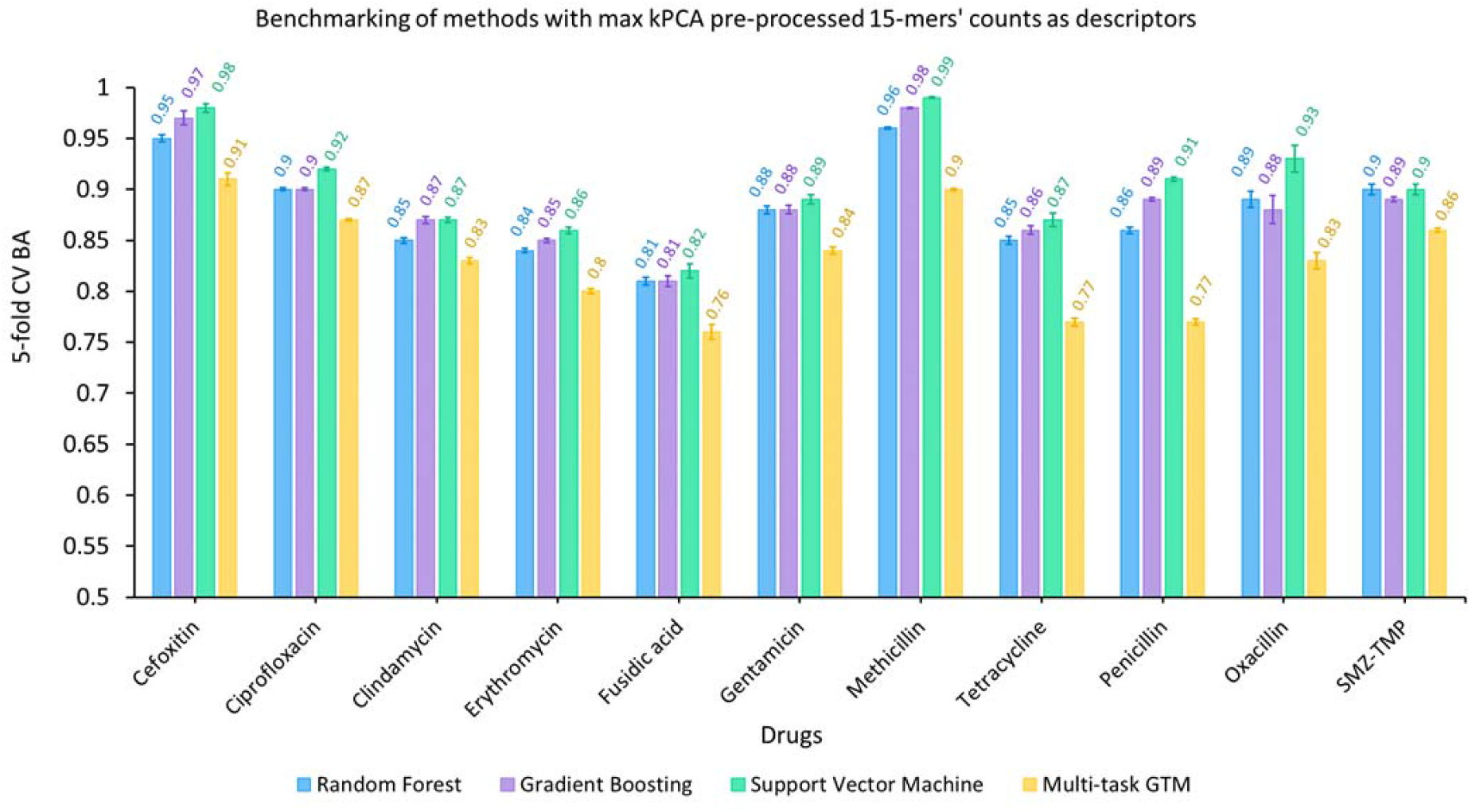
Mean values of BA obtained in 5-fold cross-validation repeated five times for multi-task GTM (yellow) and single-task Random Forest (blue), Support Vector Machine (green), Gradient Boosting (purple) for 11 antibiotics. The descriptors used for building the models were kPCA pre-processed 15-mers’ counts with ‘max’ kernel function. SMZ-TMP stands for sulfamethoxazole/trimethoprim.

Although not being among the models with the highest precision, the GTM-based models possess the advantage in the form of direct comparison of clusters of sequences in terms of resistance. In addition, AMR-gene-specific landscapes can be also combined with resistance landscapes to elucidate markers of resistance proper to some sequence clusters on the map. This provides an additional and more intuitively interpretable means for analysis of relationships between antimicrobial resistance and AMR genes. The landscapes generated for two of the antibiotics (oxacillin, sulfamethoxazole/trimethoprim) that were not used in the optimization of the models are shown together with other landscapes in Fig. S5. Clear delineation of zones related to resistant and susceptible sequences for these drugs validates that the created models can be used to predict the resistance to the drugs unseen by the model in the optimization.

### 3.2 Cartography of *S. aureus* sequence space and antibiotic resistance profiling

The herein created GTM-based resistance landscapes encompass 5773 mutant sequences of *S. aureus* isolates and are coloured according to the mean resistance status of the isolates residing in a given node of the map to the antibiotic of interest (Fig. 2). The position of each isolate (a unique genomic sequence) is not determined by two dimensions (e.g. x- and y-axes). Instead, it is defined by a probability distribution across all the nodes (i.e. squares on the map). Namely, the position of each sequence is defined by a so-called responsibility vector, whose individual values reflect the probabilities of a sequence to reside in each map node. The maps allow to easily delineate the zones containing exclusively genomes of resistant isolates (red zones) or susceptible ones (blue zones) from the zones jointly populated by both categories (colours between red and blue). White zones on the resistance landscapes correspond to the empty regions, where no sequences with drug-specific resistance profiles were found. Hence, the landscapes for drugs with the smallest number of labelled sequences contain more white zones. Saturated colours correspond to highly populated regions, while high transparency defines sparsely populated zones. Global density (technically referred to as “cumulated responsibility”) is proportional to the number of mapped sequences while the map areas covered is an expression of the relative diversity of mapped items. Both global density and diversity increase for “classical” antibiotics - methicillin, penicillin for which a lot of (and, implicitly, diverse) data is available. The highest prediction performance was attained with Jaccard kPCA preprocessing for 10- and 30-mer counts, and kPCA with ‘max’ kernel function for 15-mers. kPCA was hence the preferred pre-processing strategy. The GTMs built using k-mers with different lengths (10-, 15-, 30-mers) possessed similar predictive accuracy (Fig. S10). We have chosen the ones based on 15-mers for further analysis since they provide a more illustrative exploration of the genotype-phenotype relationships and genetic determinants of the resistance.

Equivalently to other methods aimed to analyze the genetic space and diversity of *S. aureus* isolates [19], [38], the GTM highlights the existence of several clusters of *S. aureus* strains. This is visible on all of the maps (Fig. S5). Since all of the landscapes rely on the same manifold, drug-specific landscapes can be thus directly compared. For example, the resistance of *S. aureus* isolates to •-lactam antibiotics such as penicillin and semisynthetic •-lactamase resistant penicillins, i.e. methicillin are notably different. Indeed, one can expect these differences since methicillin was developed to circumvent penicillin resistance [39]. In both landscapes, resistants (red) and susceptibles (blue) are fairly well separated on the landscapes, for both of these drugs. However, the penicillin landscape in Fig. S8 displays some more red areas (resistants) which in the neighbouring methicillin map appear as blue – these are the penicillin-resistant genomes of *S. aureus* that were successfully targeted by methicillin. However, several nodes in the right middle area and right bottom corner (Fig. S8) of the aforementioned landscapes accommodate methicillin-resistant *S. aureus* (MRSA) isolates that maintain susceptibility to penicillin, which is an example of penicillin-susceptible methicillin-resistant *S. aureus*. The incidence of such kind of resistance profile was already registered by [40]. The MRSA isolates exert resistance not only to methicillin but also to all of the other •-lactam class antibiotics, i.e. cephalosporins and carbapenems [41] although they can be susceptible to the newest classes of cephalosporins [42]. The •-lactam antibiotics such as penicillins e.g. penicillin, methicillin and cefoxitin, macrolides such as erythromycin, and quino-lone antibiotics e.g. ciprofloxacin possess landscapes with resistant clusters that are widely distributed over the entire area of the map compared to the landscapes of other antibiotics considered here (Fig. 2, Fig. S5). This testifies about a wider variety of genetic determinants that can influence the development of resistance to these three antibiotics given the current data. Macrolides such as erythromycin are usually and solely prescribed in case of trivial MSSA infections [42], [43]. This tendency is visible upon comparison of erythromycin and methicillin resistance landscapes (Fig. 2). Namely, almost all isolates susceptible to methicillin also appear to be susceptible to erythromycin, which is in accordance with literature data.

**Fig. 2.**
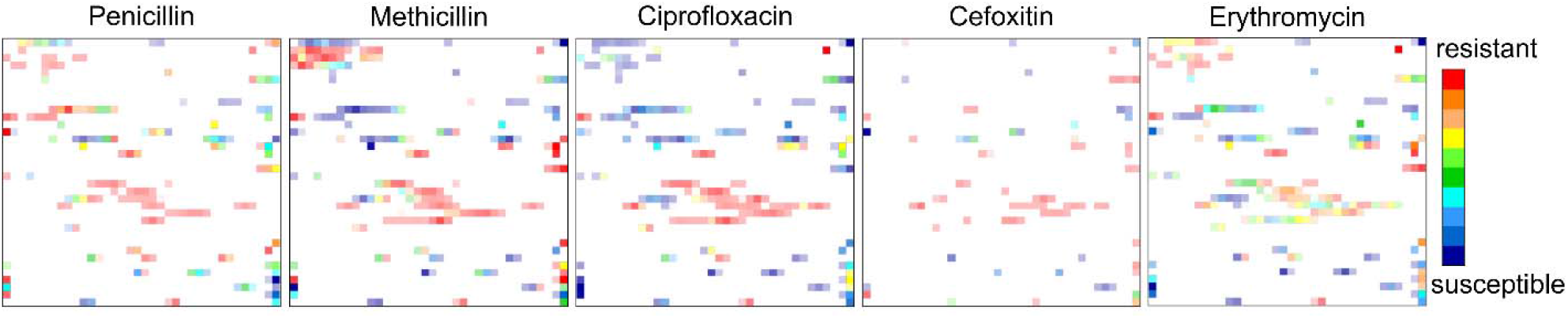
Resistance landscapes for five antibiotics against S. aureus built on kPCA pre-processed (‘max’ kernel function) 15-mers’ counts of 5773 genomic sequences. Each node is coloured by the weighted average of the antibiotic resistance profiles of the residing S. aureus isolates. Red zones are occupied by the genomes of the resistant isolates, while blue zones contain the genomes of the susceptible ones. All colours in between correspond to mixed zones containing both of them. The transparency reflects the genomes’ population density.

### 3.3 Analysis of resistance determinants using genome features landscapes

Besides the comparative analysis of the resistance profiles, the GTM can be used to analyse the distribution of the genomic features and their relationship with the resistance. For this purpose, data on genomic features, which are defined segments of genome usually related to protein and RNA-coding genes, present in the PATRIC database were used [27]. The genomes from PATRIC are annotated by these features using either the original information provided in GenBank or annotations assigned using a RAST pipeline [27], [44]. In this context, the maps can be used either to visualize the distribution of sequences containing the specific genetic feature and compare it with the resistance landscapes or select a specific part of the map and determine which genetic features are abundant in this particular zone on the map.

To show the utility of described analysis we identified the zone on drug-specific landscapes accommodating sequences with different resistance profiles to the drugs in question (Fig. 3 A). This zone of four GTM nodes contained 40 genomes of isolates conferring resistance to penicillin, but being susceptible to methicillin and other drugs. It appears in blue on the penicillin binding protein 2a (PBP2a) landscape, meaning that isolates populating this zone do not contain PBP2a and therefore it is not the reason of penicillin resistance development in them. From the analysis of other gene feature landscapes presented here, the penicillin resistance in the isolates from the selected zone can be related to, e.g. the expressed •-lactamase and/or helicase C-terminal domain (CTD) protein. The PBP2A in its turn is present in other genomes from red zones (resistant to penicillin and methicillin). Hence, the expression of PBP2a alone or in combination with other genetic determinants such as •-lactamase may be important for protecting *S. aureus* isolates from •-lactam antibiotics action such as penicillin and methicillin [45]. The PBP2a encoded by mecA gene is known to possess low affinity to •-lactam antibiotics, hence not allowing them to inhibit the bacterial cell wall biosynthesis [46]. It is worth noting, that the genomes residing in these zones contain numerous other genomic features, including, for example, genes encoding class A •-lactamases or a gene encoding helicase CTD protein (Fig. 3 B). While one cannot directly attribute the specific resistance profile based on such observations, complex genotypephenotype relationships involving the analysis of several resistance landscapes and gene features landscapes on the same map is a perspective tool for deciphering resistance profiles in the context of evolution.

**Fig. 3.**
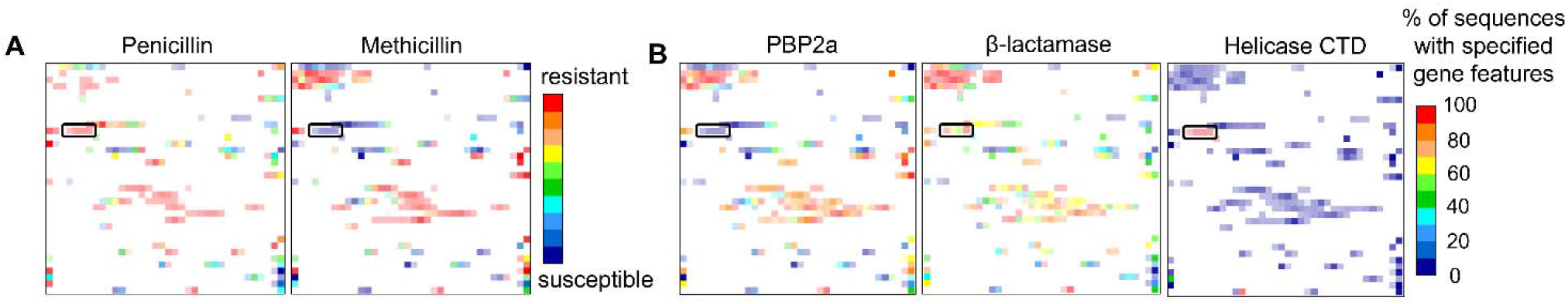
A) Resistance landscapes for penicillin and methicillin. B) Gene features’ landscapes (PBP 2A, β-lactamase, Helicase CTD) for 5773 S. aureus genomes from the frame set. Colour scheme for the resistance landscape is the same as in the Figure 2. Colour scheme for gene features landscapes reflects the percentage of genomes with specified gene features. Red zones are predominantly occupied by the genomes with specified gene features, while the blue zones contain genomes without. All colours in between correspond to mixed zones with both types of genomes in various proportions.

In addition to the observations made for the combination of the resistance landscapes and gene feature landscapes, one can use the latter to investigate the structure of the bacterial population. Horizontal gene transfer frequently occurring in bacteria leads to complex population structure: descendants of the same clone may acquire a diverse set of auxiliary genes. This complicates the analysis of the genomic space with the most common tool – phylogenetic trees. In this regard, network-based approaches and dimensionality reduction techniques represent a perspective alternative. Indeed, while some genomes comprising specific genome features resided in tight clusters (e.g. epidermin biosynthesis protein (epiC)) Fig. S13 B, others appear at various zones on the map (e.g. PBP2a, •-lactamase, helicase CTD protein, CAAX Proteases and Bacteriocin-Processing (CPBP) metalloprotease) (Fig. S13 B). In this way, the genome features landscapes can help to reveal complex evolutionary paths present in the dataset. In summary, GTM allows for illustrative exploration of the genomic space rendering the sequences with genetic determinants responsible for drug resistance development in clusters that are in accordance with the predicted resistance profiles. It can be used as a complementary tool to other methods for the analysis of genotype-phenotype relationships.

## 4 Conclusion

This paper introduces a methodology for genomic space exploration and genotype-phenotype modelling in the context of AMR prediction. This methodology allows one to represent complex genomic data in the form of interpretable 2D maps serving both as a visualization tool and a predictive ML model. It can be used as a complementary tool to guide clinical decisions in the early stages of diagnostics to provide a timely antibiotic treatment all along allowing healthcare professionals to intuitively understand the results of predictions. Herein, the introduced methodology was tested for the analysis of *S. aureus* genome space, but it is readily amenable to other bacteria. Furthermore, the suggested approach is big data-compatible and is able, in principle, to accumulate an unlimited amount of data. At the same time, it is also capable to operate well with limited quantity of data for building models. In this context, GTM can be used as a tool for visualization and analysis of large bacterial genomic data being complementary to standard tools such as phylogenetic trees and network-based approaches.

## Acknowledgment

Corresponding author: Alexandre Varnek

## Conflict of interest

none declared.

The data underlying this article are available at https://doi.org/10.5281/zenodo.7101559.

## Supplementary information

### Text S1

The data pre-processing protocol consisted in the following steps applied to the data extracted from PATRIC:

1. Selection of genomes with the AMR phenotypes labelled as “Susceptible”/“Resistant”. Intermediate and ambiguous labels such as “Not defined”, “IS”, “r”, “RS”, “Susceptible dose-dependent”, etc. were excluded.
2. Removal of genome duplicates with ambiguous resistance profile, namely with the presence of both susceptibility and resistance to the same drug.
3. The labels “0”/“1” were assigned to “Susceptible”/“Resistant” genomes respectively.
4. The AMR profiles for the same drug combinations named differently in PATRIC (e.g. sulfamethoxazole/trimethoprim and sulfamethoxazole-trimethoprim) were merged to increase the number of data points.
5. The drugs, possessing AMR data only for one of the classes (susceptible or resistant labels only) were removed from the dataset.
6. The drugs that contained less than 45 data points in either of the AMR classes and less than 100 data points overall for each of the bacteria species were removed from the dataset.
7. The drugs with highly imbalanced AMR records distribution, namely, the “0” to “1” or “1” to “0” class ratio higher than 3.5 were subjected to undersampling. The undersampling was carried out by decreasing the majority class size, i.e. by randomly removing data points of the majority class until reaching the class ratio of 1:1.

**Table S1.** The number of susceptible and resistant data points to 11 antibiotics against S.aureus used in this study.

|  | **Antibiotics** | **Resistant** | **Susceptible** |
| --- | --- | --- | --- |
| *S.aureus* | Cefoxitin | 247 | 247 |
|  | Ciprofloxacin | 1436 | 2834 |
|  | Clindamycin | 1100 | 2111 |
|  | Erythromycin | 1952 | 2678 |
|  | Fusidic acid | 332 | 332 |
|  | Gentamicin | 686 | 686 |
|  | Methicillin | 2315 | 1845 |
|  | Tetracycline | 643 | 643 |
|  | Penicillin | 2519 | 919 |
|  | Oxacillin | 91 | 91 |
|  | Sulfamethoxazole/trimethoprim | 361 | 361 |

### Text S2

For the generation of unitigs, the DBGWAS tool [1] was used with S.aureus DNA sequences in a form of contigs. The unitigs are representative DNA sequences of varying length that were obtained via compacted De Bruijin graphs (DBGs) construction from k-mers (the process of unitigs generation is shown in Figure S3). In the current work, for the generation of unitigs the k-mers of length 31 were used since previous benchmarking studies confirmed the optimal performance of 31-mers for genomes comparison [2] among others (k-mers of length 15, 21, 51, 71, 91). The overall number of generated 31-mers for 5773 annotated genomic sequences of S.aureus was 48.8 M, from which the overall number of generated unitigs was 2.4 M. The descriptors were pre-processed in a way to remove rare and quasi-constant descriptors, hence leading to the reduction of the feature space to ~976 K (number of unitigs). Quasi-constant (variance threshold of 0.05) and rare descriptors (k-mers present in •1 % of the dataset) were filtered out since their inability to differentiate across sequences. However, such significant feature space dimensionality reduction was still not sufficient, thus compelling us to apply Jaccard kernel PCA (kPCA) pre-processing and finally retrieve first 300 principal components (PCs) as descriptors. The 300 first PCs were chosen as they allowed us to preserve the variance of the original dataset.

### Text S3

PCA was performed using scikit-learn library v. 1.1.1. kPCA was performed using scikit-learn library v. 1.1.1. The number of components was kept to 300, while similarity matrices (Jaccard, max, min ani, cosine, Mash) generated by Kmer-db were used as kernels.

### Text S4

Generative topographic mapping (GTM), first proposed by Bishop [3], is one of the most efficient dimensionality reduction methods. It performs non-linear projections of objects from the initial multidimensional descriptor space to a 2D latent space - a manifold defined by a set of radial basis functions (RBF). The shape and position of each point of the man ifold in the N-dimensional space are determined during its training – unsupervised fitting to the “frameset” items - objects used to probe the sequence space of interest. Afterward, the manifold is unfolded back to the planar form – square grid 2D map (Figure S1).

Once trained, the manifold can host not only the objects of the “frameset” but also any external objects, under the condition that in the multidimensional space, they are residing close to the manifold (log Likelihood control [4]). The distinctive feature and the main advantage of GTM is its probabilistic nature, ensured by RBFs. In GTM, projected objects are not assigned to a particular point on the map. Instead, each one is fuzzily projected over the whole map with larger probabilities (“responsibilities”) for nodes situated closer to this object in the initial space. Such smooth projection enables the creation of GTM landscapes – 2D plots of cumulated responsibilities, coloured by average values of different properties, e. g. density, biological activity, etc. One manifold can host multiple landscapes allowing the analysis of multiple libraries according to different properties.

Detailed description of the GTM algorithm is given below.

**Figure S1.**
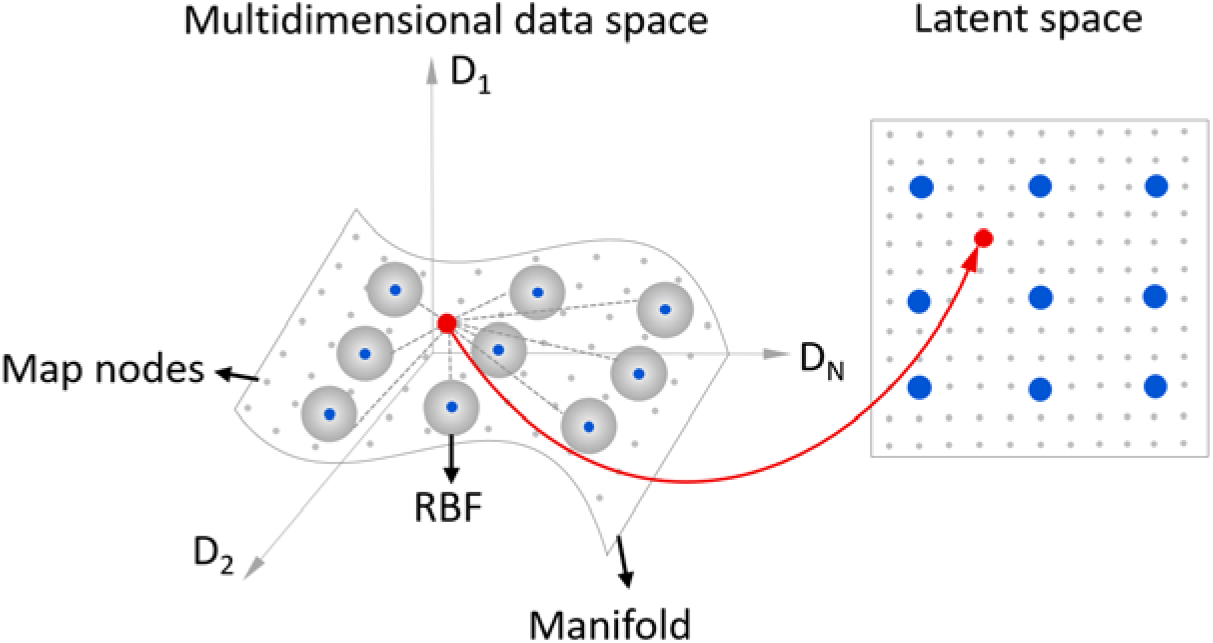
The data point describing the sequence in multidimensional space is projected in the 2D latent space.

At the first step of a GTM construction, a bounded flexible 2D hypersurface called manifold defined by a linear combination of Gaussian Radial Basis Functions (RBFs) is injected into the multidimensional space of the sequences. The coordinates of the manifold in the multidimensional space (embedding the sequences in our case) are obtained using the Y mapping function – an inner product of • and W matrices:

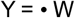

Where W is the parameter matrix defined as a (W=M×D) with M – the number of RBFs, D – dimensionality of the initial high-dimensional space (the number of sequence descriptors used) and • defined as (•K×M) with K - the number of nodes in 2D latent space.

The • consists of RBF estimations on each node of the manifold, whereas W consists of weight values describing the manifold in the multidimensional space. The training of the manifold implies optimization of the manifold to reach the densest regions of frame set data space (a set representative protein sequences). To train the manifold, firstly the W is initialized. This is done by fitting the manifold to the plane defined by the two first principal components from a PCA of the frame set. Subsequently, the initialized manifold is bent in the D-dimensional space to approach the data points as close as possible. The fitting is done using Expectation-Maximization (EM) algorithm. The target is to maximize the (natural logarithm of the) probability density *p*(*t_n_*|*x_k_*, *W, β*) of a sequence *t_n_* considering a normal distribution of width *β*, centered on the manifold *W* evaluated at each node *x_k_*: the (log-)likelihood or Llh (1(2).

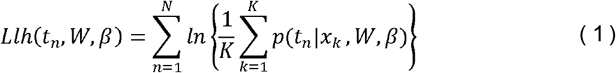

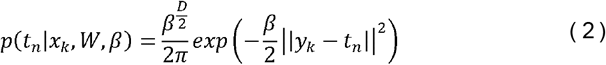

Where

*Llh*(*t_n_, W, β*) – log-likelihood function of the sequence

*p*(*t_n_*|*x_k_*, *W*, *β*) – probability density of the sequence in multidimensional space to be associated with the node in 2D la tent grid

– point representing a sequence in the multidimensional descriptor space
  coordinates of a node in the 2D latent space
  coordinates of a node in the multidimensional space
  common inverse variance of the distribution
- parameter matrix

Once the model is trained, the next step consists in a projection of sequences from the frame set onto the manifold. Using the Bayes formula, the responsibilities are computed at each node and for each sequence.

Therefore, each sequence projected onto the manifold possesses Llh value and responsibility vector (vector of values each designating the probability of a sequence to reside in a particular node). The Llh value reflects the optimal closeness of the sequence to the manifold, whereas the responsibility vector describes the sequence as a probability distribution in the 2D latent space. The sum of responsibilities for each of the map nodes allows one to analyse the distribution of sequences over the map and can be visualized as a density landscape. (Figure S2 A)

To build a property landscape, the probability of a sequence labelled as resistant or susceptible to belong to a node is defined as following (3):

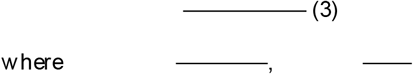

- number of sequences associated with the class, – total number of sequences used for training, – sum of sequences responsibilities in the class, for the node k. Each node is colored by the weighted average of class ratios of residing sequences yielding a classification landscape (Figure S2 B).

The prediction of the class for the new sequence is performed as following:

It should be noted that the same manifold can accommodate several property landscapes. Since the training of the manifold is unsupervised, the models based on the activity landscapes may perform less good than some state of the art supervised machine learning approaches.

**Figure S2.**
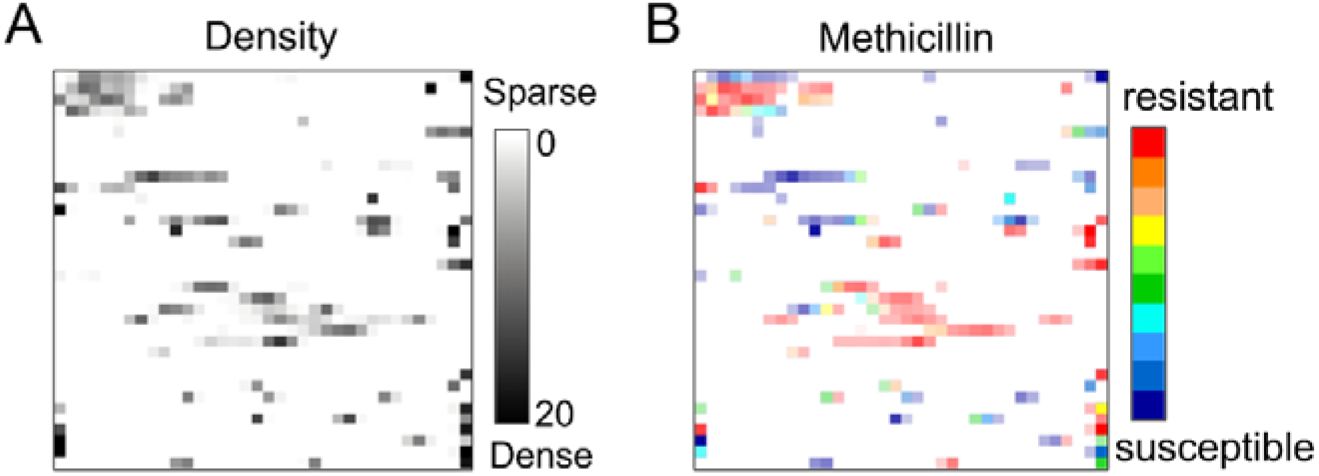
A) Density landscape; B) Class landscape.

**Figure S3.**
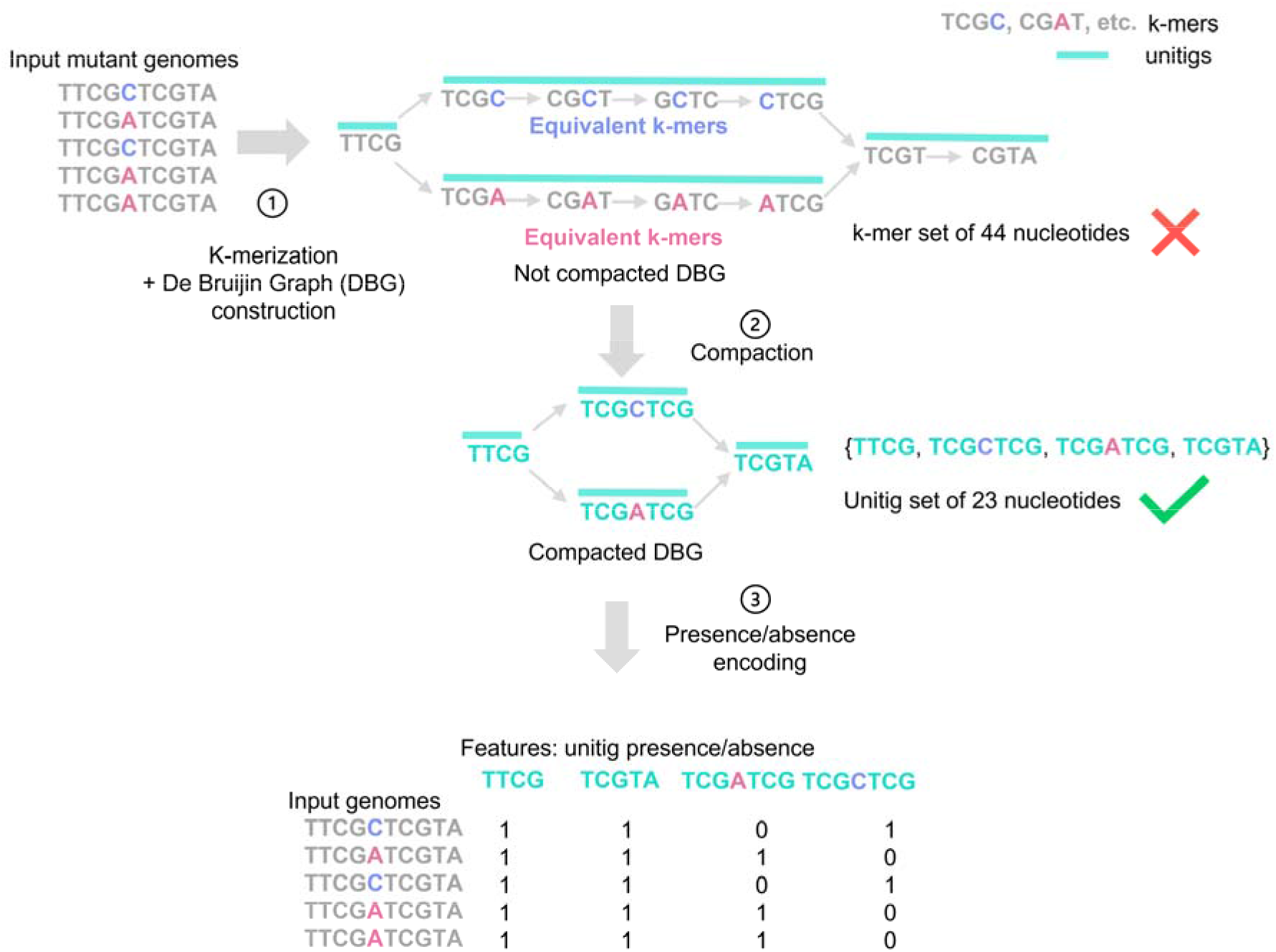
Unitigs generation process. More details on unitigs generation can be found in the paper by Jaillard et al. [1].

### Text S5

The following hyperparameters were tuned during optimization (grid search):

- RF [5]: number of trees (100, 300, 500, 1000), number of features (squared root of the number of features, log2 of number of features), out-of-bag sampling (with and without), max depth of trees (full tree, 5, 10, 30), class weight (none, balanced, balanced_subsample);
- SVM [6]: regularization coefficient (0.1, 1, 10, 100, 1000), kernel coefficient (1, 0.1, 0.01, 0.001, 0.0001), kernel (‘rbf’, ‘linear’, ‘poly’, ‘sigmoid’);
- GB (Friedman, 2001): number of trees (100, 300, 500, 1000), number of features (squared root of the number of features, log2 of number of features), learning rate (0.0001, 0.001, 0.01, 0.1, 1.0), subsampling (0.5, 0.7, 1.0) max depth of trees (full tree, 5, 10, 30).

**Figure S4.**
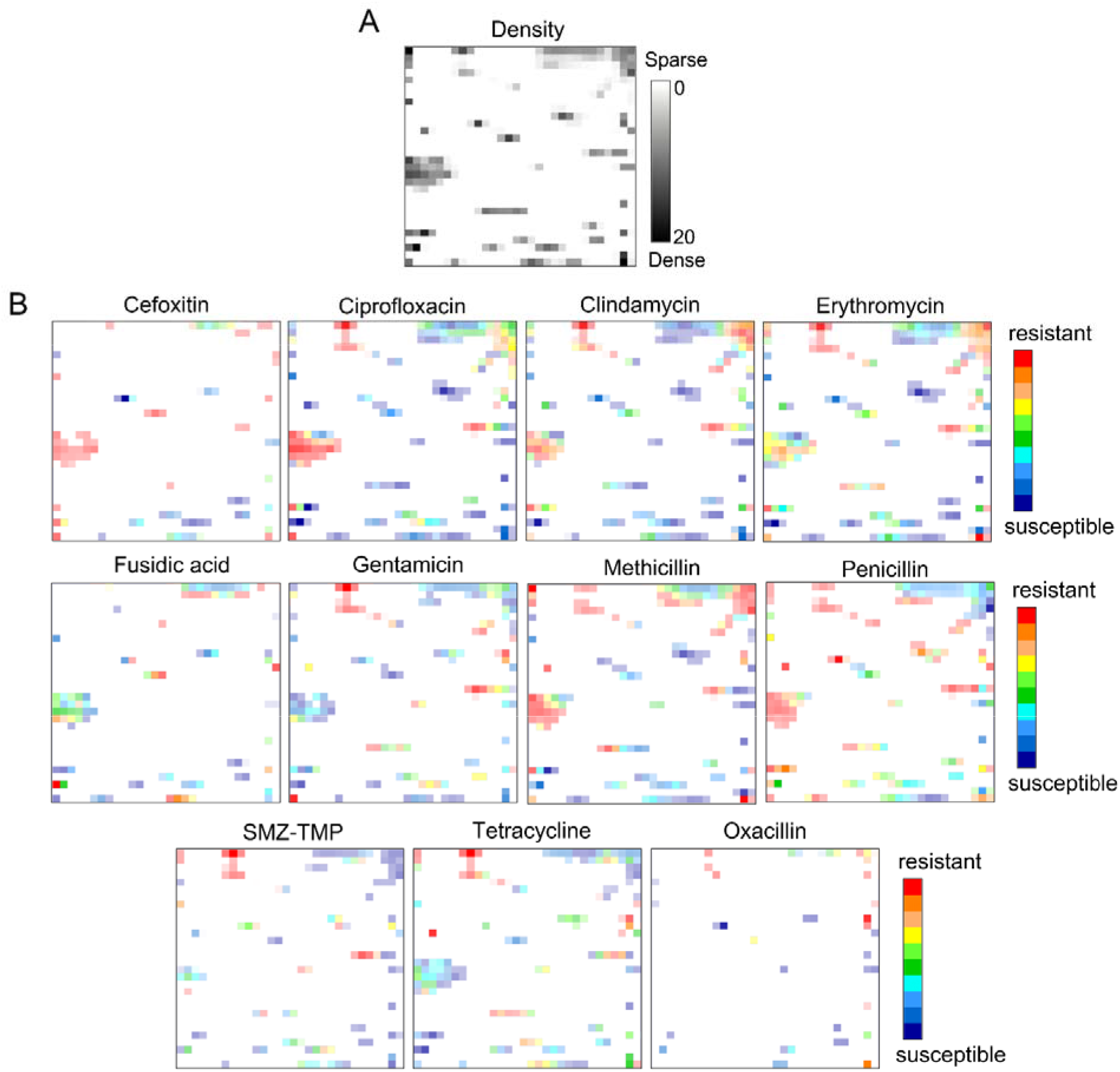
Resistance landscapes for 11 antibiotics against S.aureus built on Jaccard kPCA pre-processed 10-mers’ counts of 5773 sequences. Each node is coloured by the weighted average of the of antibiotic resistance profiles of the residing S.aureus isolates. Red zones are occupied by the genomes of the resistant isolates, while the blue zones contain genomes of susceptible ones. All colours in between correspond to mixed zones containing both of them. The transparency reflects thegenomes’ population density. SMZ-TMP stands for sulfamethoxazole/trimethoprim antibiotic combination.

**Figure S5.**
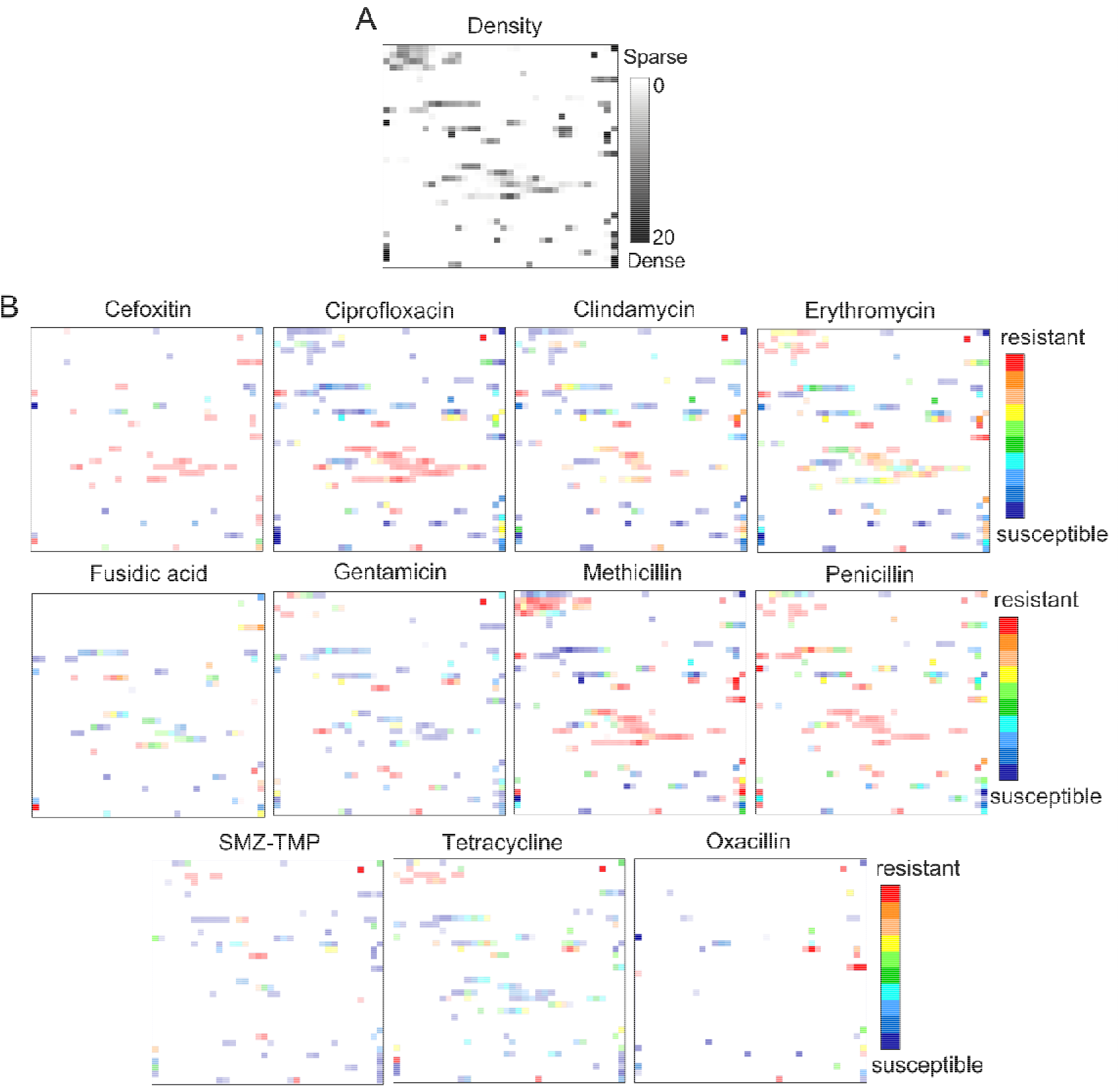
Resistance landscapes for 11 antibiotics against S.aureus built on max kPCA pre-processed 15-mers’ counts of 5773 sequences. Each node is coloured by the weighted average of the of antibiotic resistance profiles of the residing S.aureus isolates. Red zones are occupied by the genomes of the resistant isolates, while the blue zones contain the genomes of the susceptible ones. All colours in between correspond to mixed zones containing both of them. The transparency reflects the genomes’ population density. SMZ-TMP stands for sulfamethoxazole/trimethoprim antibiotic combination.

**Figure S6.**
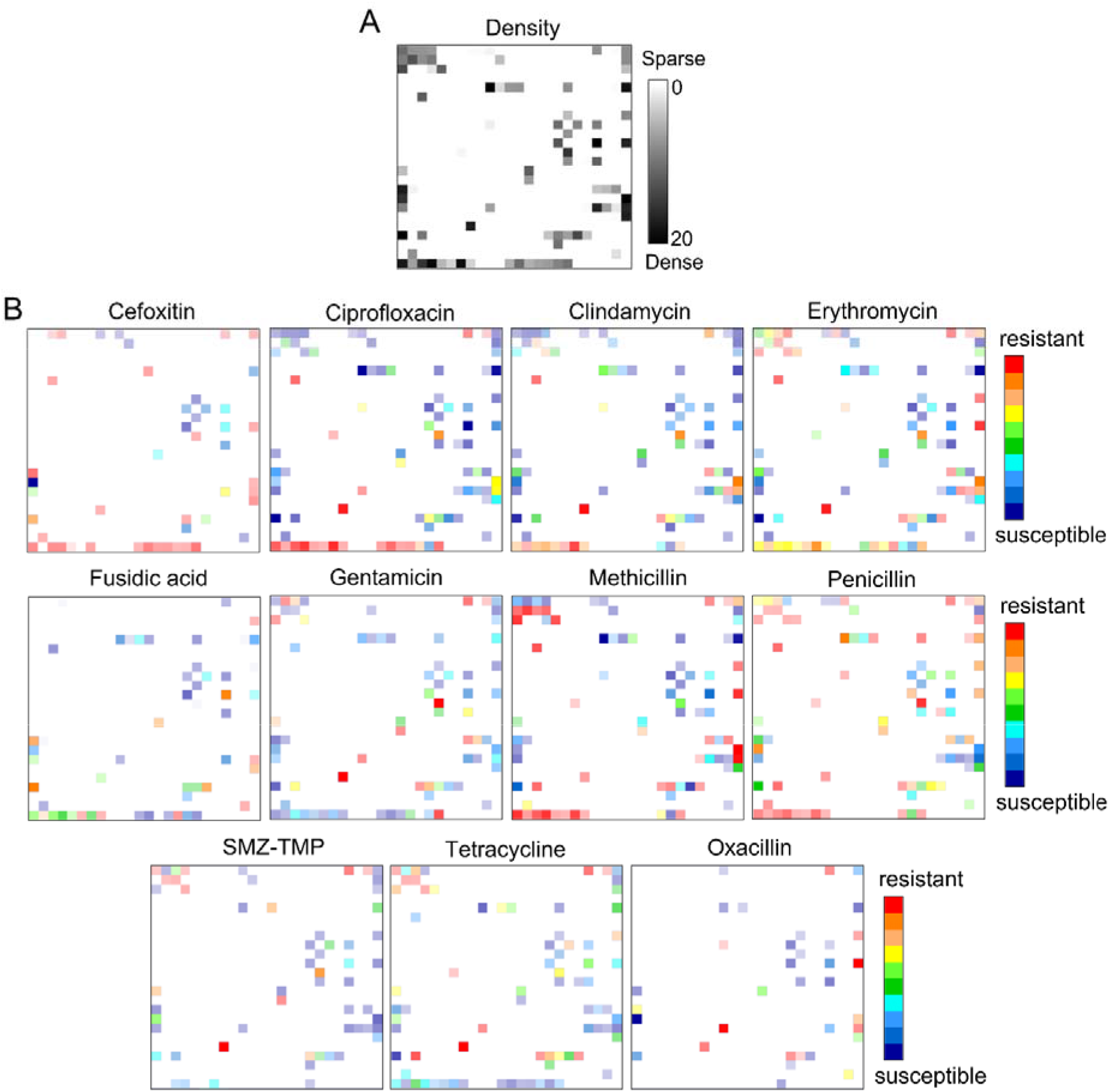
Resistance landscapes for 11 antibiotics against S.aureus built on Jaccard kPCA pre-processed 30-mers’ counts of 5773 sequences. Each node is coloured by the weighted average of the of antibiotic resistance profiles of the residing S.aureus isolates. Red zones are occupied by the genomes of the resistant isolates, while the blue zones contain the genomes of susceptible ones. All colours in between correspond to mixed zones containing both of them. The transparency reflects the genomes’ population density. SMZ-TMP stands for sulfamethoxazole/trimethoprim antibiotic combination.

**Figure S7.**
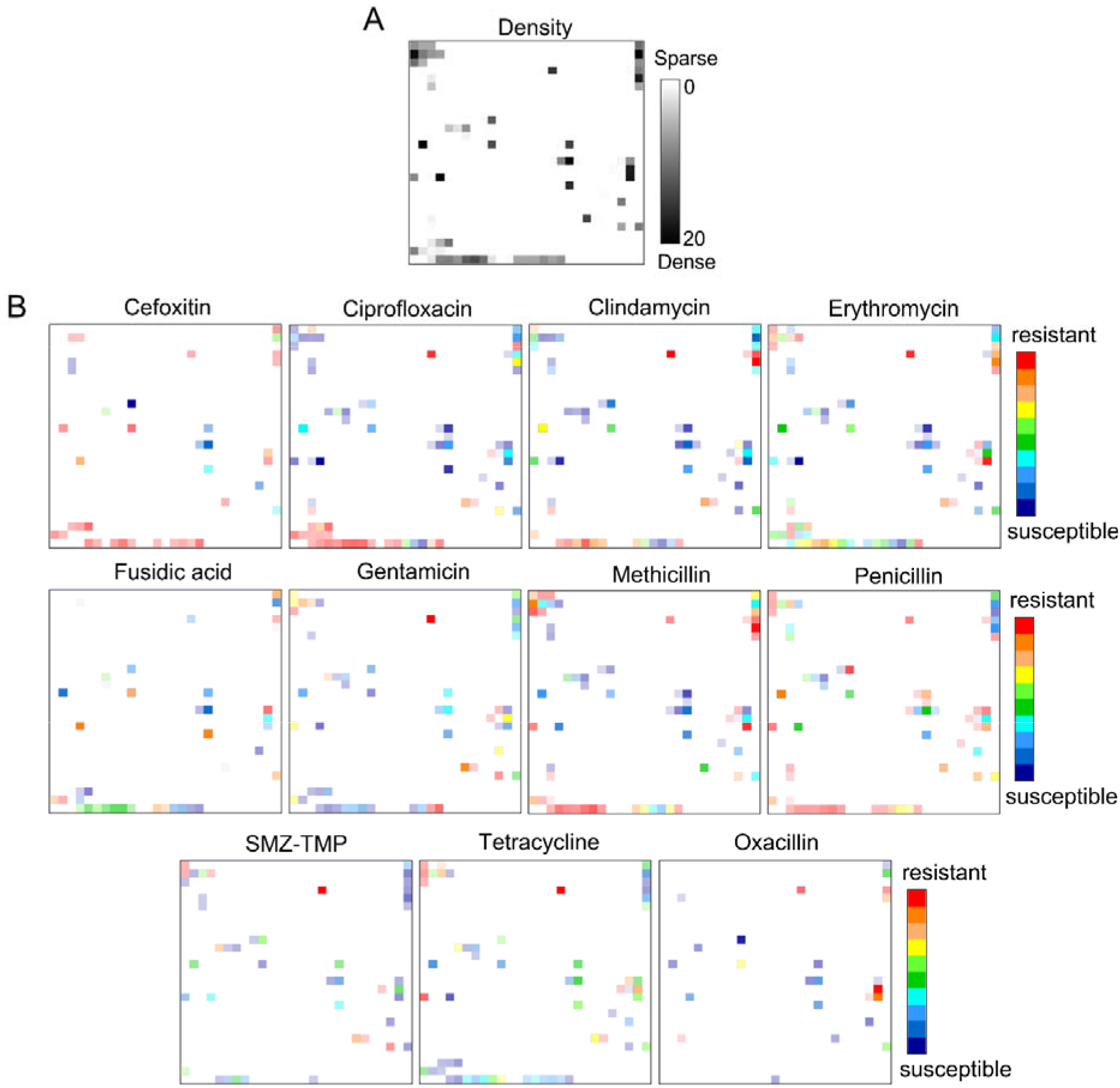
Resistance landscapes for 11 antibiotics against S.aureus built on Jaccard kPCA pre-processed unitigs’ presence/absence vectors of 5773 sequences. Each node is coloured by the weighted average of the of antibiotic resistance profiles of the residing S.aureus isolates. Red zones are occupied by the genomes of the resistant isolates, while the blue zones contain the genomes of susceptible ones. All colours in between correspond to mixed zones containing both of them. The transparency reflects the genomes’ population density. SMZ-TMP stands for sulfamethoxazole/trimethoprim antibiotic combination.

**Figure S8.**
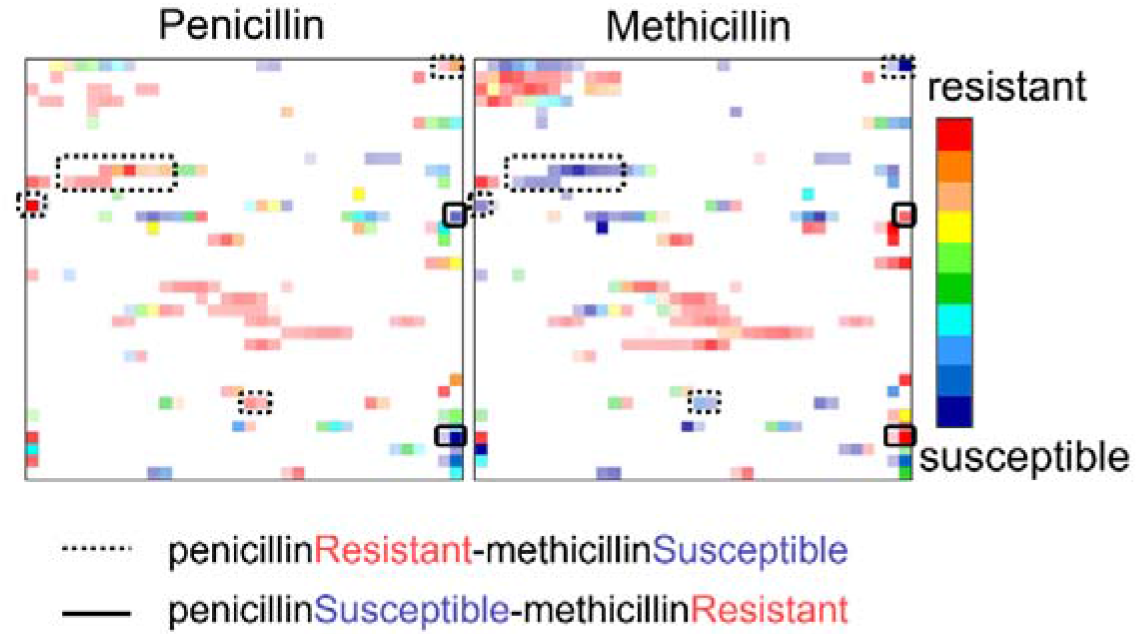
Areas in resistance landscapes marked by black continuous rectangles are populated by MRSA isolates that maintain susceptibility to penicillin, whereas rectangles in dashed line are populated by penicillin-resistant genomes of S. aureus that were successfully targeted by methicillin.

**Figure S9.**
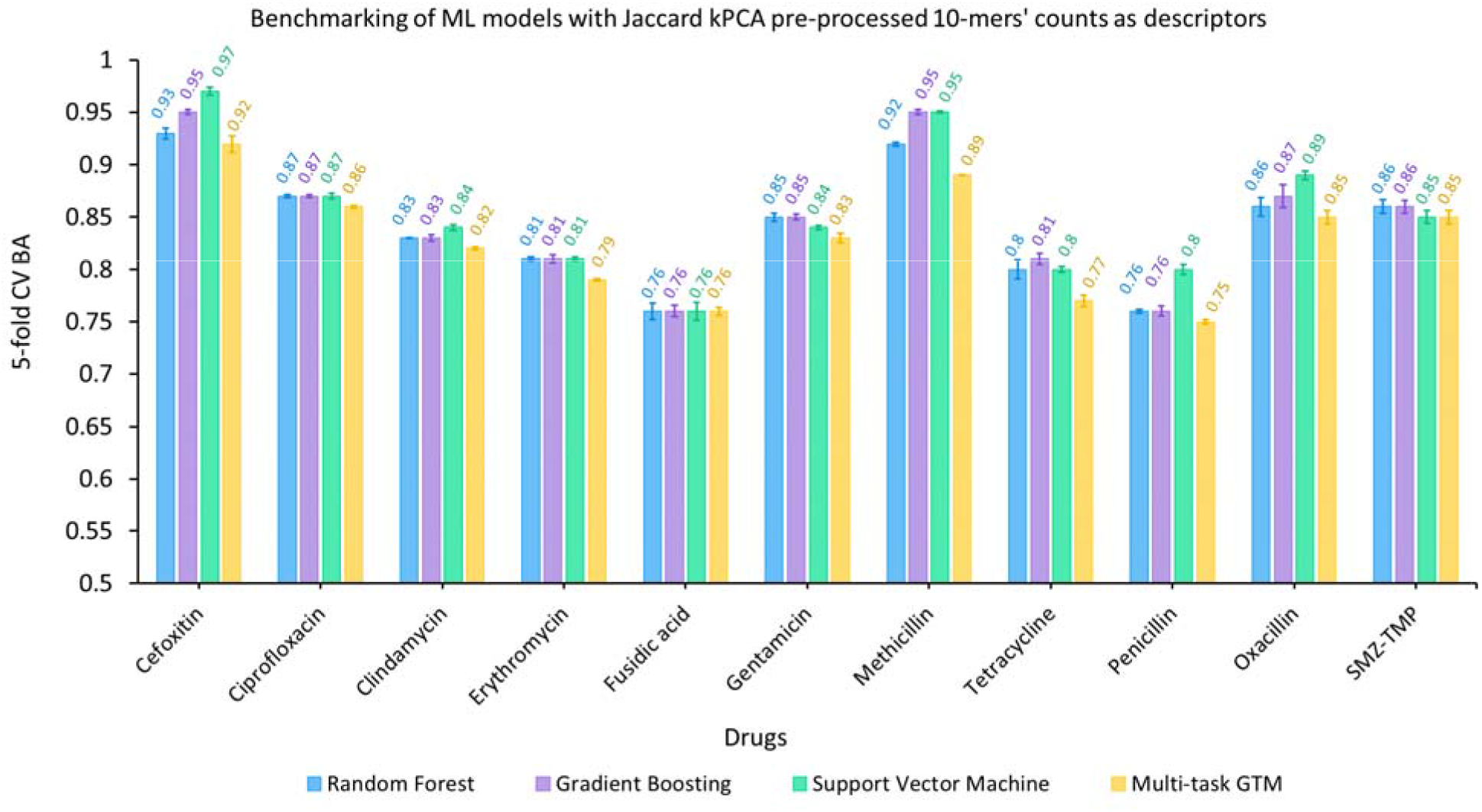
Mean values of BA obtained in 5-fold cross-validation repeated five times for multi-task GTM (yellow) and single task Random Forest (blue), Support Vector Machine (green), Gradient Boosting (purple) for 11 antibiotics. The descriptors used for building the models were Jaccard kPCA pre-processed 10-mers’ counts.

**Figure S10.**
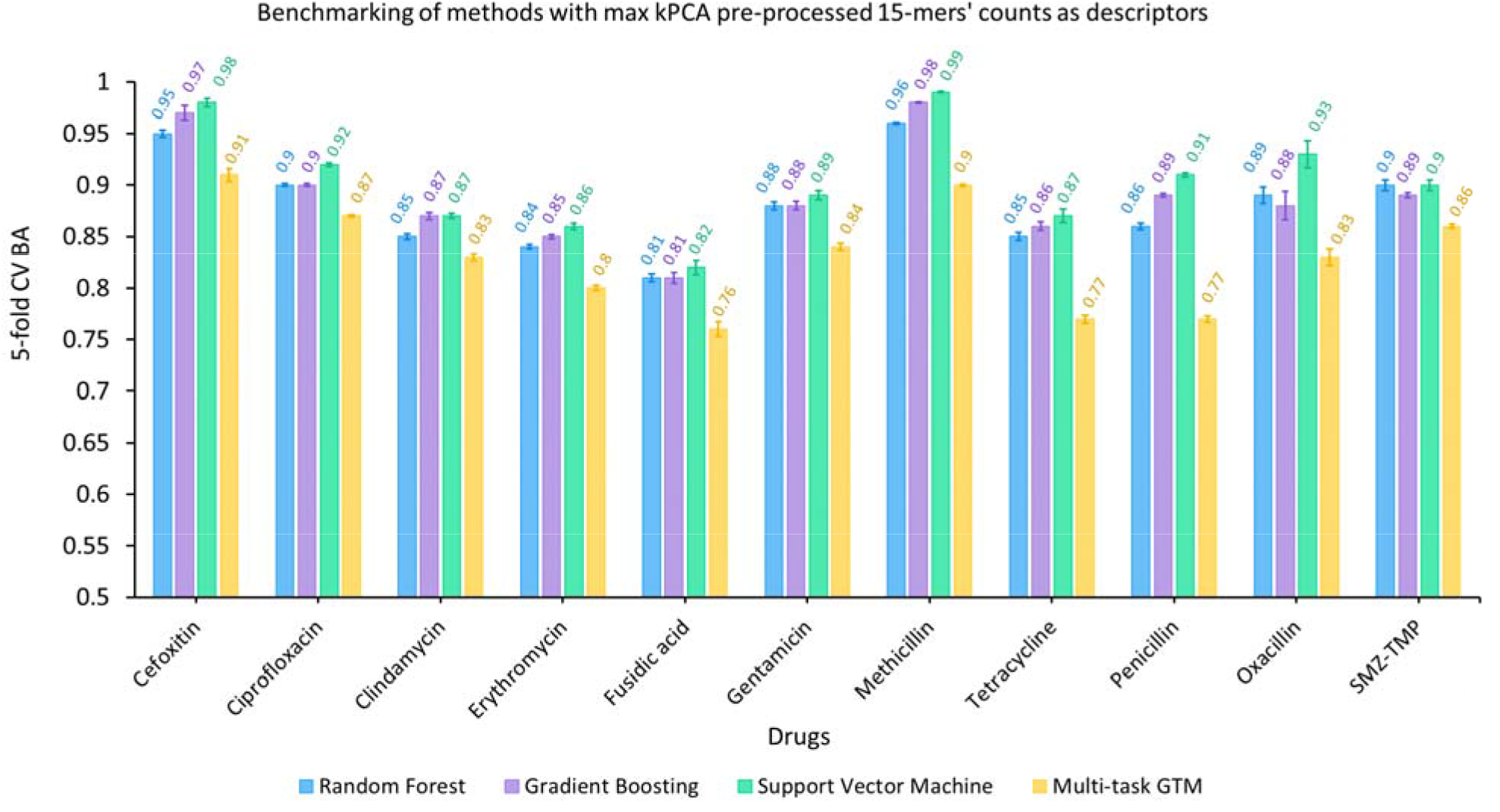
Mean values of BA obtained in 5-fold cross-validation repeated five times for multi-task GTM (yellow) and single task Random Forest (blue), Support Vector Machine (green), Gradient Boosting (purple) for 11 antibiotics. The descriptors used for building the models were kPCA pre-processed 15-mers’ counts with ‘max’ kernel function.

**Figure S11.**
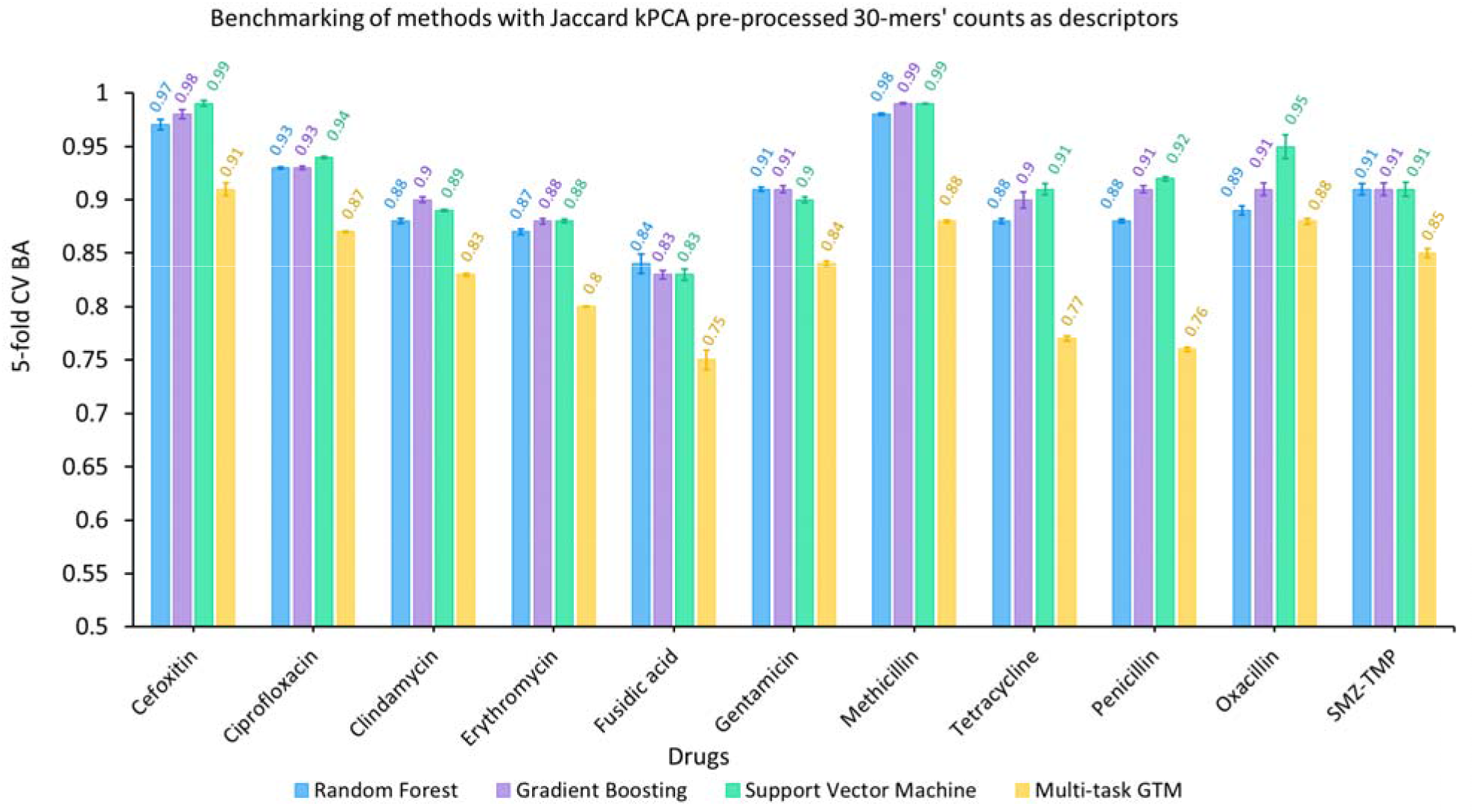
Mean values of BA obtained in 5-fold cross-validation repeated five times for multi-task GTM (yellow) and single task Random Forest (blue), Support Vector Machine (green), Gradient Boosting (purple) for 11 antibiotics. The descriptors used for building the models were Jaccard kPCA pre-processed 30-mers’ counts.

**Figure S12.**
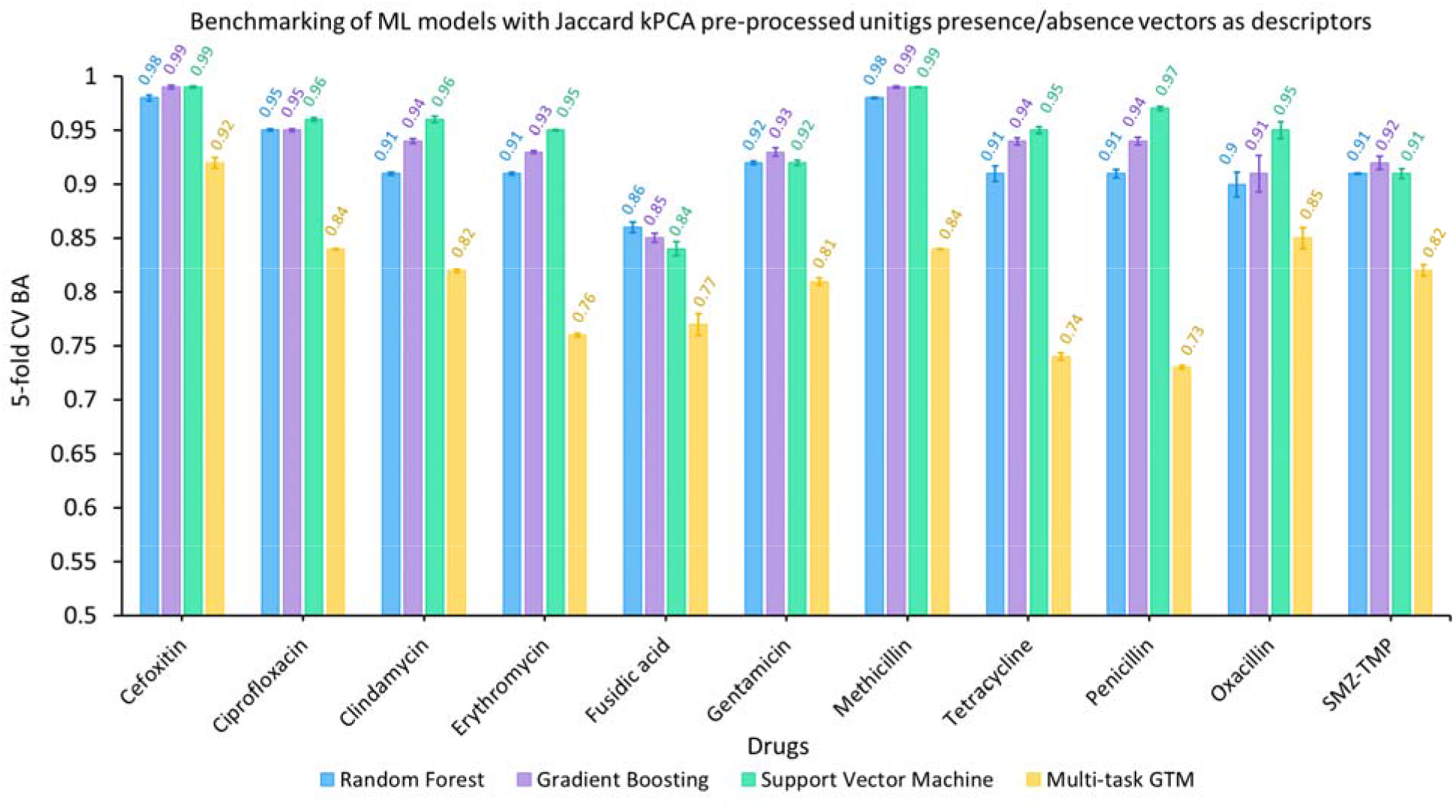
Mean values of BA obtained in 5-fold cross-validation repeated five times for multi-task GTM (yellow) and single task Random Forest (blue), Support Vector Machine (green), Gradient Boosting (purple) for 11 antibiotics. The descriptors used for building the models were Jaccard kPCA pre-processed unitigs’ presence/absence vectors.

**Figure S13.**
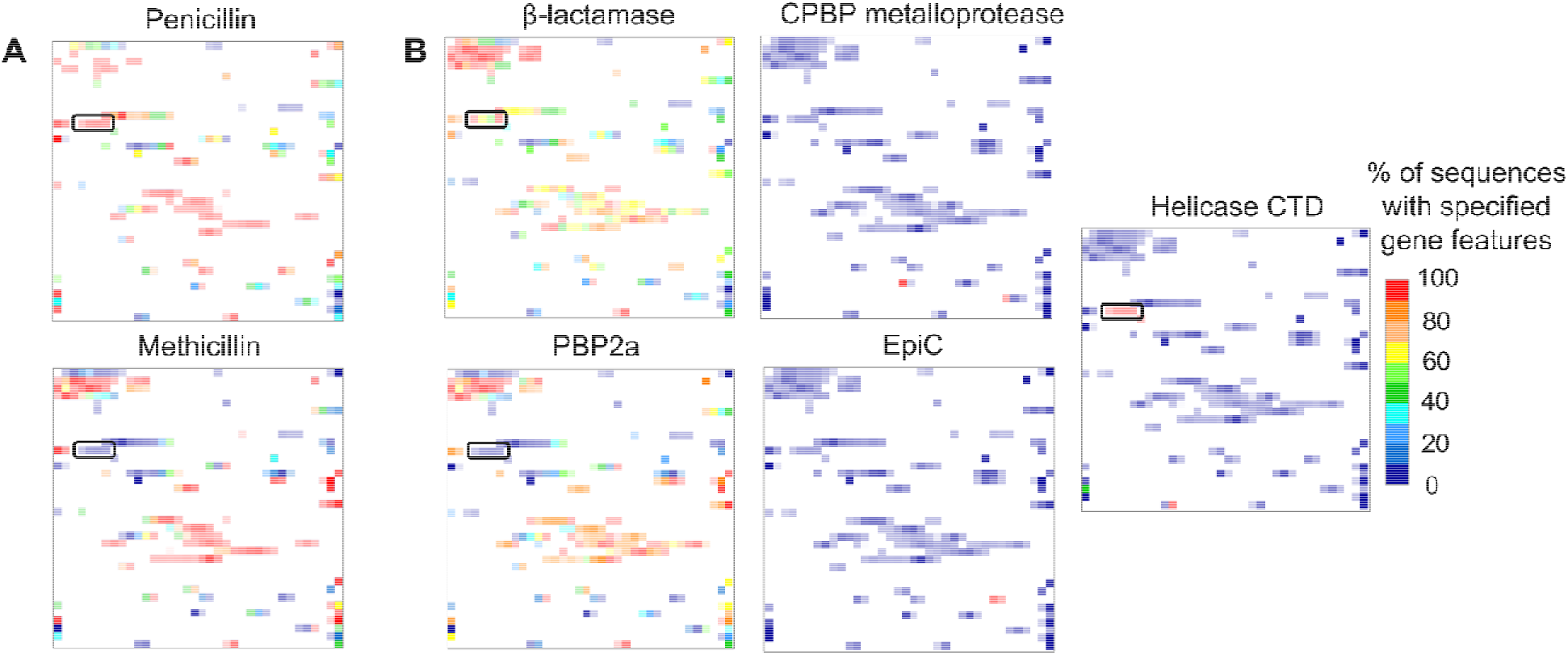
A) Resistance landscapes for penicillin and methicillin. B) Gene features’ landscapes (PBP 2A, β-lactamase, CPBP metalloprotease, EpiC, Helicase CTD) for 5773 S.aureus genomes from the frame set. Color scheme for the resistance landscape is the same as in the Figure S5. Color scheme for gene features landscapes reflects the percentage of genomes with specified gene features. Red zones are predominantly occupied by the genomes with specified gene features, while the blue zones contain genomes without. All colors in between correspond to mixed zones containing both types of genomes in various proportions.

## Notes

https://doi.org/10.5281/zenodo.7101559

### Competing Interest Statement

The authors have declared no competing interest.

## References

[1] C. J. Antimicrobial Resistance Collaborators et al., “Global burden of bacterial antimicrobial resistance in 2019: a systematic analysis.,” Lancet (London, England), vol. 399, no. 10325, pp. 629–655, Feb. 2022, doi: 10.1016/S0140-6736(21)02724-0.

[2] J. O’Neill, “Tackling drug-resistant infections globally: Final report and recommendations. The review on antimicrobial resistance.,” 2016, [Online]. Available: https://amr-review.org/sites/default/files/160525_Finalpaper_withcover.pdf.

[3] M. Anderson et al., “Averting the AMR crisis□: What are the avenues for policy?,” WHO Reg. Off. Eur. - Heal. Syst. policy Anal., pp. 1–34, 2019.

[4] J. H. Jorgensen and M. J. Ferraro, “Antimicrobial susceptibility testing: A review of general principles and contemporary practices,” Clin. Infect. Dis., vol. 49, no. 11, pp. 1749–1755, 2009, doi: 10.1086/647952.

[5] A. van Belkum et al., “Developmental roadmap for antimicrobial susceptibility testing systems,” Nat. Rev. Microbiol., vol. 17, no. 1, pp. 51–62, 2019, doi: 10.1038/s41579-018-0098-9.

[6] M. N. Anahtar, J. H. Yang, and S. Kanjilal, “Applications of Machine Learning to the Problem of Antimicrobial Resistance: an Emerging Model for Translational Research,” J. Clin. Microbiol., vol. 59, no. 7, Jul. 2021, doi: 10.1128/JCM.01260-20.

[7] P.-J. Van Camp, D. B. Haslam, and A. Porollo, “Bioinformatics Approaches to the Understanding of Molecular Mechanisms in Antimicrobial Resistance.,” Int. J. Mol. Sci., vol. 21, no. 4, Feb. 2020, doi: 10.3390/ijms21041363.

[8] J. J. Davis et al., “Antimicrobial Resistance Prediction in PATRIC and RAST.,” Sci. Rep., vol. 6, p. 27930, Jun. 2016, doi: 10.1038/srep27930.

[9] M. Yasir, A. Mustafa, S. Kausar, A. A. Bajaffer, and E. I. Azhar, “Saudi Journal of Biological Sciences Prediction of antimicrobial minimal inhibitory concentrations for Neisseria gonorrhoeae using machine learning models,” Saudi J. Biol. Sci., vol. 29, no. 5, pp. 3687–3693, 2022, doi: 10.1016/j.sjbs.2022.02.047.

[10] D. Moradigaravand, M. Palm, A. Farewell, V. Mustonen, J. Warringer, and L. Parts, “Prediction of antibiotic resistance in Escherichia coli from large-scale pan-genome data,” PLoS Comput. Biol., vol. 14, no. 12, p. e1006258, Dec. 2018, doi: 10.1371/journal.pcbi.1006258.

[11] Y. Ren et al., “Prediction of antimicrobial resistancea based on whole-genome sequencing and machine learning,” Bioinformatics, vol. 38, no. 2, pp. 325–334, Jan. 2022, doi: 10.1093/bioinformatics/btab681.

[12] D. Aytan-Aktug, P. T. L. C. Clausen, V. Bortolaia, F. M. Aarestrup, and O. Lund, “Prediction of Acquired Antimicrobial Resistance for Multiple Bacterial Species Using Neural Networks,” mSystems, vol. 5, no. 1, Jul. 2020, doi: 10.1128/mSystems.00774-19.

[13] O. Bonham-Carter, J. Steele, and D. Bastola, “Alignment-free genetic sequence comparisons: a review of recent approaches by word analysis,” Brief. Bioinform., vol. 15, no. 6, pp. 890–905, Nov. 2014, doi: 10.1093/bib/bbt052.

[14] A. Drouin, G. Letarte, F. Raymond, M. Marchand, J. Corbeil, and F. Laviolette, “Interpretable genotype-to-phenotype classifiers with performance guarantees,” Sci. Rep., vol. 9, no. 1, p. 4071, Jul. 2019, doi: 10.1038/s41598-019-40561-2.

[15] M. Jaillard, M. Palmieri, A. van Belkum, and P. Mahé, “Interpreting k-mer–based signatures for antibiotic resistance prediction,” Gigascience, vol. 9, no. 10, p. giaa110, Oct. 2020, doi: 10.1093/gigascience/giaa110.

[16] H. A. Gaspar and G. Breen, “Probabilistic ancestry maps: a method to assess and visualize population substructures in genetics,” BMC Bioinformatics, vol. 20, no. 1, p. 116, 2019, doi: 10.1186/s12859-019-2680-1.

[17] J. A. Lees, G. Tonkin-Hill, Z. Yang, and J. Corander, “Mandrake: visualising microbial population structure by embedding millions of genomes into a low-dimensional representation,” bioRxiv, p. 2021.10.28.466232, 2021, [Online]. Available: https://www.biorxiv.org/content/10.1101/2021.10.28.466232v1 https://www.biorxiv.org/content/10.1101/2021.10.28.466232v1.abstract.

[18] S. Sakaue et al., “Dimensionality reduction reveals fine-scale structure in the Japanese population with consequences for polygenic risk prediction,” Nat. Commun., vol. 11, no. 1, p. 1569, 2020, doi: 10.1038/s41467-020-15194-z.

[19] D. C. Melles et al., “Natural population dynamics and expansion of pathogenic clones of Staphylococcus aureus.,” J. Clin. Invest., vol. 114, no. 12, pp. 1732–1740, Dec. 2004, doi: 10.1172/JCI23083.

[20] M. Nicolau, A. J. Levine, and G. Carlsson, “Topology based data analysis identifies a subgroup of breast cancers with a unique mutational profile and excellent survival,” Proc. Natl. Acad. Sci., vol. 108, no. 17, pp. 7265–7270, Apr. 2011, doi: 10.1073/pnas.1102826108.

[21] P. G. Cámara, “Topological methods for genomics: present and future directions.,” Curr. Opin. Syst. Biol., vol. 1, pp. 95–101, Feb. 2017, doi: 10.1016/j.coisb.2016.12.007.

[22] C. M. Bishop, M. Svensén, and C. K. I. Williams, “GTM: The Generative Topographic Mapping,” 1998.

[23] N. Kireeva, I. I. Baskin, H. A. Gaspar, D. Horvath, G. Marcou, and A. Varnek, “Generative Topographic Mapping (GTM): Universal Tool for Data Visualization, Structure-Activity Modeling and Dataset Comparison,” Mol. Inform., vol. 31, no. 3–4, pp. 301–312, 2012, doi: https://doi.org/10.1002/minf.201100163.

[24] D. Horvath, G. Marcou, and A. Varnek, “Generative topographic mapping in drug design,” Drug Discov. Today Technol., vol. 32–33, pp. 99–107, 2019, doi: 10.1016/j.ddtec.2020.06.003.

[25] K. Pikalyova et al., “HIV-1 drug resistance profiling using amino acid sequence space cartography,” Bioinformatics, p. btac090, Feb. 2022, doi: 10.1093/bioinformatics/btac090.

[26] N. R. Wray et al., “Genome-wide association analyses identify 44 risk variants and refine the genetic architecture of major depression,” Nat. Genet., vol. 50, no. 5, pp. 668–681, 2018, doi: 10.1038/s41588-018-0090-3.

[27] J. J. Davis et al., “The PATRIC Bioinformatics Resource Center: expanding data and analysis capabilities.,” Nucleic Acids Res., vol. 48, no. D1, pp. D606–D612, Jan. 2020, doi: 10.1093/nar/gkz943.

[28] G. Rizk, D. Lavenier, and R. Chikhi, “DSK: k-mer counting with very low memory usage,” Bioinformatics, vol. 29, no. 5, pp. 652–653, Mar. 2013, doi: 10.1093/bioinformatics/btt020.

[29] S. Deorowicz, A. Gudy•, M. Długosz, M. Kokot, and A. Danek, “Kmer-db: instant evolutionary distance estimation,” Bioinformatics, vol. 35, no. 1, pp. 133–136, Jan. 2019, doi: 10.1093/bioinformatics/bty610.

[30] T. Kohonen, “The Self-Organizing Map,” Proc. IEEE, vol. 78, no. 9, pp. 1464–1480, 1990, doi: 10.1109/5.58325.

[31] L. Van Der Maaten and G. Hinton, “Visualizing Data using t-SNE,” 2008.

[32] D. Horvath, J. Brown, G. Marcou, and A. Varnek, “An Evolutionary Optimizer of libsvm Models,” Challenges, vol. 5, no. 2, pp. 450–472, 2014, doi: 10.3390/challe5020450.

[33] A. Lin, D. Horvath, G. Marcou, B. Beck, and A. Varnek, “Multi-task generative topographic mapping in virtual screening.,” J. Comput. Aided. Mol. Des., vol. 33, no. 3, pp. 331–343, Mar. 2019, doi: 10.1007/s10822-019-00188-x.

[34] L. Breiman, “Random Forests,” Mach. Learn., vol. 45, no. 1, pp. 5–32, 2001, doi: 10.1023/A:1010933404324.

[35] B. E. Boser, I. M. Guyon, and V. N. Vapnik, “A Training Algorithm for Optimal Margin Classifiers,” in Proceedings of the Fifth Annual Workshop on Computational Learning Theory, 1992, pp. 144–152, doi: 10.1145/130385.130401.

[36] J. H. Friedman, “Greedy function approximation: A gradient boosting machine.,” Ann. Stat., vol. 29, no. 5, pp. 1189–1232, 2001, doi: 10.1214/aos/1013203451.

[37] F. Pedregosa et al., “Scikit-learn: Machine Learning in Python,” J. Mach. Learn. Res., vol. 12, no. 85, pp. 2825–2830, 2011, [Online]. Available: http://jmlr.org/papers/v12/pedregosa11a.html.

[38] H. F. L. Wertheim et al., “The role of nasal carriage in Staphylococcus aureus infections.,” Lancet. Infect. Dis., vol. 5, no. 12, pp. 751–762, Dec. 2005, doi: 10.1016/S1473-3099(05)70295-4.

[39] R. Knox, “A new penicillin (BRL 1241) active against penicillin-resistant staphylococci.,” Br. Med. J., vol. 2, no. 5200, pp. 690–693, Sep. 1960, doi: 10.1136/bmj.2.5200.690.

[40] D. S. Blanc et al., “Unusual spread of a penicillin-susceptible methicillin-resistant Staphylococcus aureus clone in a geographic area of low incidence.,” Clin. Infect. Dis. an Off. Publ. Infect. Dis. Soc. Am., vol. 29, no. 6, pp. 1512–1518, Dec. 1999, doi: 10.1086/313522.

[41] A. Pantosti and M. Venditti, “What is MRSA?,” Eur. Respir. J., vol. 34, no. 5, pp. 1190–1196, 2009, doi: 10.1183/09031936.00007709.

[42] T. J. Foster, “Antibiotic resistance in Staphylococcus aureus. Current status and future prospects.,” FEMS Microbiol. Rev., vol. 41, no. 3, pp. 430–449, May 2017, doi: 10.1093/femsre/fux007.

[43] C. Rayner and W. J. Munckhof, “Antibiotics currently used in the treatment of infections caused by Staphylococcus aureus.,” Intern. Med. J., vol. 35 Suppl 2, pp. S3–16, Dec. 2005, doi: 10.1111/j.1444-0903.2005.00976.x.

[44] T. Brettin et al., “RASTtk: a modular and extensible implementation of the RAST algorithm for building custom annotation pipelines and annotating batches of genomes.,” Sci. Rep., vol. 5, p. 8365, Feb. 2015, doi: 10.1038/srep08365.

[45] M. Miragaia, “Factors contributing to the evolution of Meca-mediated •-lactam resistance in staphylococci: Update and new insights from whole genome sequencing (WGS),” Front. Microbiol., vol. 9, no. NOV, pp. 1–16, 2018, doi: 10.3389/fmicb.2018.02723.

[46] M.-A. W. Shalaby, E. M. E. Dokla, R. A. T. Serya, and K. A. M. Abouzid, “Penicillin binding protein 2a: An overview and a medicinal chemistry perspective.,” Eur. J. Med. Chem., vol. 199, p. 112312, Aug. 2020, doi: 10.1016/j.ejmech.2020.112312.

## References

[1] M. Jaillard et al., “A fast and agnostic method for bacterial genome-wide association studies: Bridging the gap between k-mers and genetic events.,” PLoS Genet., vol. 14, no. 11, p. e1007758, Nov. 2018, doi: 10.1371/journal.pgen.1007758.

[2] A. Drouin et al., “Predictive computational phenotyping and biomarker discovery using reference-free genome comparisons,” BMC Genomics, vol. 17, no. 1, p. 754, 2016, doi: 10.1186/s12864-016-2889-6.

[3] C. M. Bishop, M. Svensén, and C. K. I. Williams, “GTM: The Generative Topographic Mapping,” 1998.

[4] H. A. Gaspar, I. I. Baskin, G. Marcou, D. Horvath, and A. Varnek, “GTM-Based QSAR Models and Their Applicability Domains.,” Mol. Inform., vol. 34, no. 6–7, pp. 348–356, Jun. 2015, doi: 10.1002/minf.201400153.

[5] L. Breiman, “Random Forests,” Mach. Learn., vol. 45, no. 1, pp. 5–32, 2001, doi: 10.1023/A:1010933404324.

[6] B. E. Boser, I. M. Guyon, and V. N. Vapnik, “A Training Algorithm for Optimal Margin Classifiers,” in Proceedings of the Fifth Annual Workshop on Computational Learning Theory, 1992, pp. 144–152, doi: 10.1145/130385.130401.

